## Supplemental Material for "Structure-guided mutagenesis of OSCAs reveals differential activation to mechanical stimuli"

### Figures

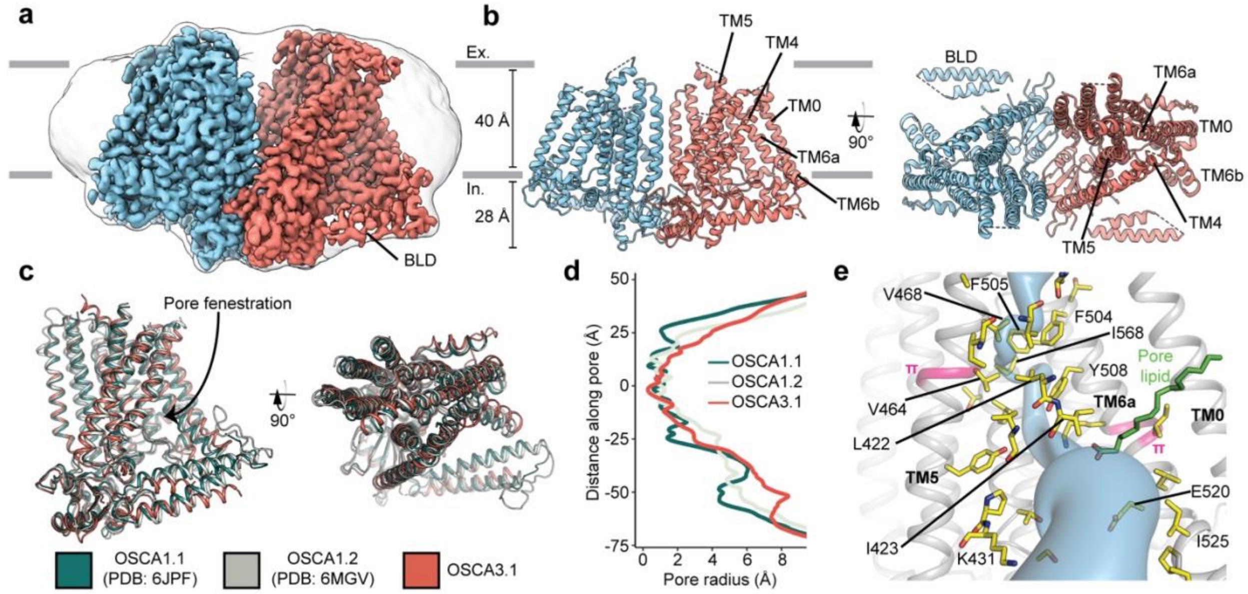


**Figure 1: Cryo-EM structure and pore of OSCA3.1.**

**a,** Cryo-EM map of OSCA3.1 dimer colored by subunit. Nanodisc density in grey corresponds to the unsharpened map (gaussian-filtered to 1.5 σ). **b,** Front (left) and top (right) view of atomic model. Ex.: Extracellular, In.: Intracellular. **c,** Superposition of OSCA1.1, OSCA1.2 (in nanodiscs) and OSCA3.1 protomers. **d,** Pore profile of OSCAs in **c**. **e,** View of the pore pathway (blue) of OSCA3.1. Pore facing residues colored in yellow, with selected residues labeled. π-helical turns in pink. Putative pore lipid in green. Backbone of TM3 and TM4 helices hidden for clarity.

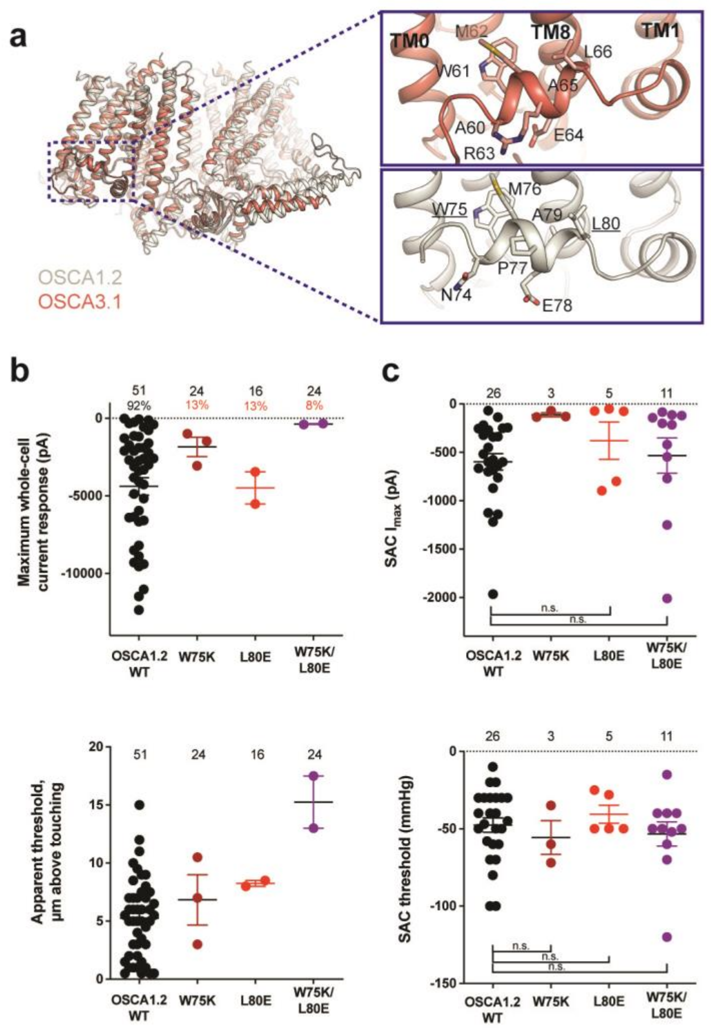


**Figure 2: Amphipathic helix mutants of OSCA1.2.**
**a,** Superposition of OSCA1.2 (grey) and OSCA3.1 (red). Insets: close-up view of amphipathic helix. Residues substituted in OSCA1.2 for electrophysiology experiments are underlined. **b,** Few cells expressing mutations in the amphipathic helix respond to poking compared to WT controls. Top panel: Maximum poke-induced currents observed in whole-cell mode for cells exposed to displacements up to 8.9 ± 0.6 µm (N=51; mean ± S.E.M.), 12.8 ± 0.7 µm (N=24), 12.4 ± 0.7 µm (N=16), and 12.8 ± 0.5 µm (N=24) above touching for WT, OSCA1.2$_{W75K}$, OSCA1.2$_{L80E}$, and OSCA1.2$_{W75K/L80E}$, respectively. Bottom panel: the apparent threshold in µm above touching the cell for this cohort. **c,** SAC maximal current ($I_{max}$) (top) and mmHg threshold (bottom) from WT and OSCA1.2$_{W75K/L80E}$-expressing cells reveal no significant differences in the ability of negative pressure to activate channels in cell-attached patches (Student's *t*-test). Also shown are data from single mutants OSCA1.2$_{W75K}$ and OSCA1.2$_{L80E}$. Too few patches were obtained for OSCA1.2$_{W75K}$ to compare $I_{max}$. Electrode resistances for **c** were similar in all cases (WT: 2.3 ± 0.1 MΩ (N=26); W75K: 2.1 ± 0.4 MΩ (N=3), L80E: 2.2 ± 0.1 MΩ (N=5), W75K/L80E: 1.8 ± 0.1 MΩ (N=11)). For panels **b-c**: individual cells are represented as scatter points; mean and S.E.M. are displayed.

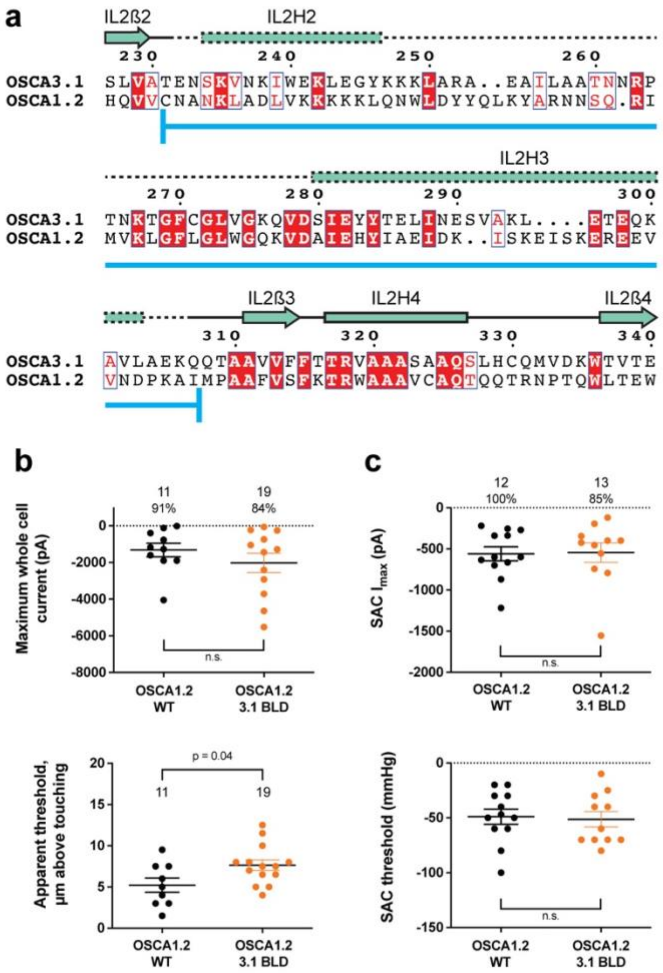


**Figure 3: OSCA1.2$_{\mathrm{OSCA3.1\text{-}BLD}}$ chimera.**

**a,** Amino acid sequence alignment of the BLD region of OSCA1.2 and OSCA3.1. Full alignment in **Supplementary Fig. 3.** Blue line at bottom of sequences denotes the sequence swapped in OSCA1.2$_{\mathrm{OSCA3.1\text{-}BLD}}$ chimera. **b,** Poke-induced responses are observed in HEK-P1KO cells expressing OSCA1.2$_{\mathrm{3.1\text{-}BLD}}$. Top panel: Maximum poke-induced currents observed in whole-cell mode for cells exposed to displacements up to 9.8 ± 0.8 µm (N=11; mean ± S.E.M.) and 11.8 ± 0.6 µm (N=19) above touching for WT and OSCA1.2$_{\mathrm{3.1\text{-}BLD}}$, respectively. The percentage of cells with responses are shown above. Bottom panel: the apparent threshold in µm above touching the cell for this cohort. **c,** SAC $I_{max}$ (top) and mmHg threshold (bottom) from WT and OSCA1.2$_{\mathrm{3.1BLD}}$-expressing cells reveal similar activity induced by negative pressure in cell-attached patches. Data shown were obtained from the same experiments. Electrode resistances for **c** were similar in all cases (WT: 2.4 ± 0.1 MΩ (N=12); OSCA1.2$_{\mathrm{OSCA3.1\text{-}BLD}}$: 2.5 ± 0.1 MΩ (N=13)).

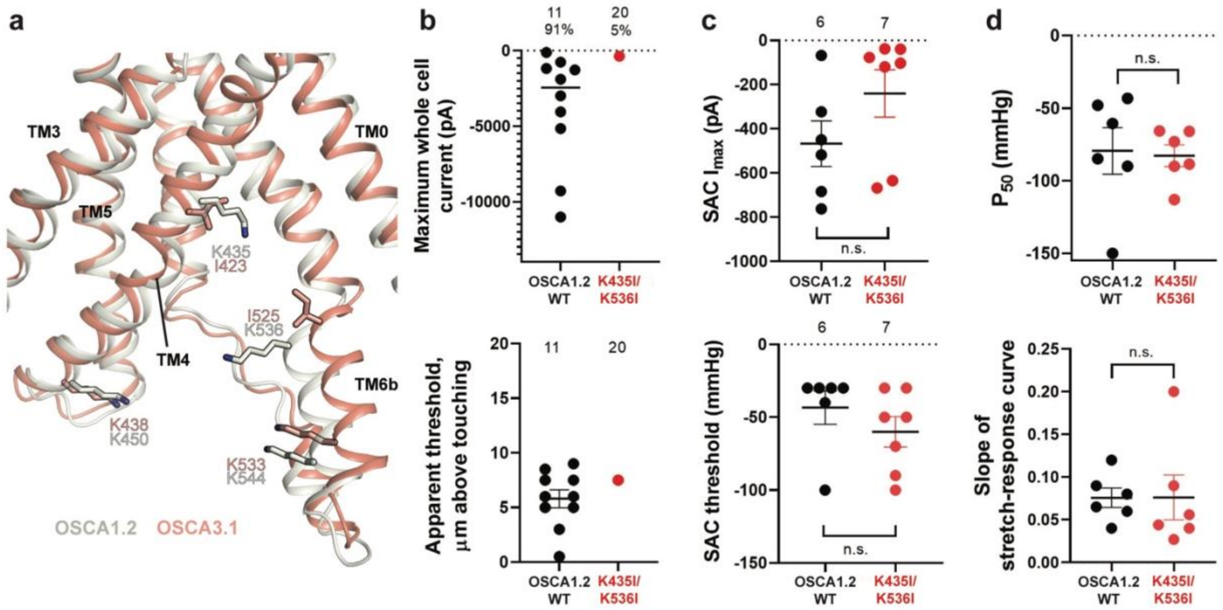


**Figure 4: Probing the functional role of the potential lipid-interacting residues.** **a,** Superposition of OSCA1.2 and OSCA3.1 around the pore fenestration. Residues of OSCA1.2 predicted to interact with lipids and corresponding residues in OSCA3.1 are shown. **b,** Few OSCA1.2$_{K435I/K536I}$ -expressing HEK-P1KO cells respond to the poke stimulus while nearly all cells expressing OSCA1.2 WT channels respond (percentages shown above under the N of cells tested). Top panel: Maximum poke-induced currents observed in whole-cell mode for cells exposed to displacements up to 9.7 ± 1 µm (N=11; mean ± S.E.M.) and 12.1 ± 0.6 µm (N=20) above touching for WT and OSCA1.2$_{K435I/K536I}$, respectively. Bottom panel: the apparent threshold in µm above touching the cell for this cohort. **c,** SAC Imax (top) and mmHg threshold (bottom) from WT and OSCA1.2$_{K435I/K536I}$-expressing cells reveal no significant differences in the ability of negative pressure to activate channels in cell-attached patches (Student's *t*-test). Electrode resistances were similar in all cases (WT: 2.1 ± 0.2 MΩ (N=10); K435I/K546I: 1.7 ± 0.3 MΩ (N=7)). Data were collected within 15 min of exposure of cells to high K+ used in this assay. **d,** Analysis of stimulus-response relationships reveal no significant differences (Student's *t*-test) in pressure to half-maximal activation ($P_{50}$; top) or the slope of the curve (bottom). Whole cell and SAC data were obtained from cells transfected at the same time.

### Supplemental figures:

#### Structure-guided mutagenesis of OSCAs reveals differential activation to mechanical stimuli

Sebastian Jojoa-Cruz[1], Adrienne E. Dubin[2], Wen-Hsin Lee[1], Andrew Ward[1]†

Affiliations

[1] Department of Integrative Structural and Computational Biology, Scripps Research, La Jolla, California 92037, USA.

[2] Department of Neuroscience, Scripps Research, La Jolla, California 92037, USA.

†Corresponding author

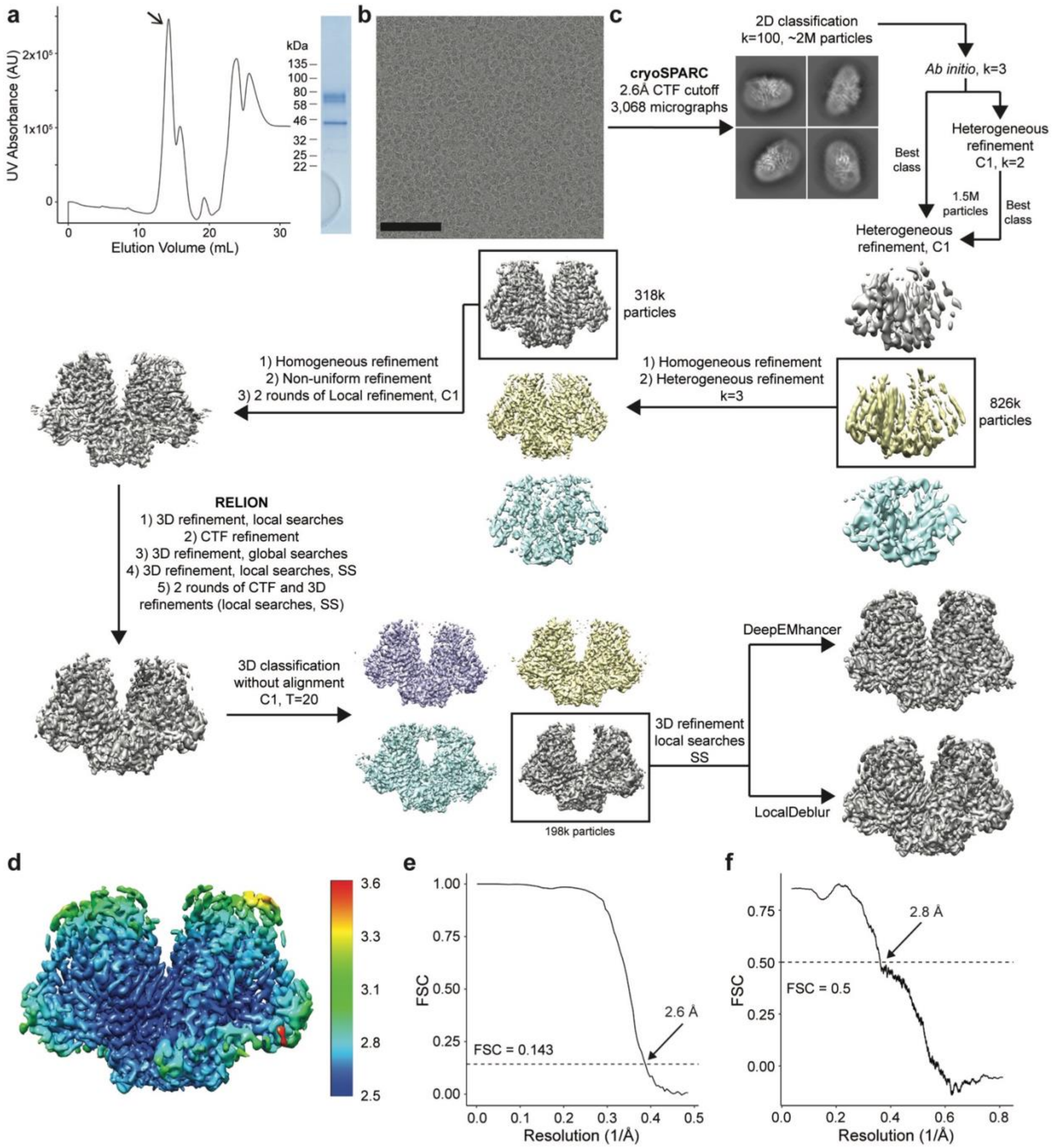


**Supplementary Figure 1: Purification and cryo-EM data processing of OSCA3.1.**
**a,** SEC trace (left) from OSCA3.1 reconstitution in nanodiscs. Arrow points to peak corresponding to fractions pooled and ran on SDS-PAGE (right). Molecular weight of OSCA3.1 and MSP2N2 is approximately 82kDa and 43kDa, respectively. **b,** Representative cryo-EM micrograph. Black bar is 100nm. **c,** Cryo-EM processing workflow. C2 symmetry was imposed during refinements unless otherwise specified. 3D refinements in RELION were performed using the SIDESPLITTER algorithm where indicated (SS). **d,** LocalDeblur map colored by local resolution calculated in RELION. **e,** FSC plot calculated in RELION. **f,** LocalDeblur map to model FSC plot.

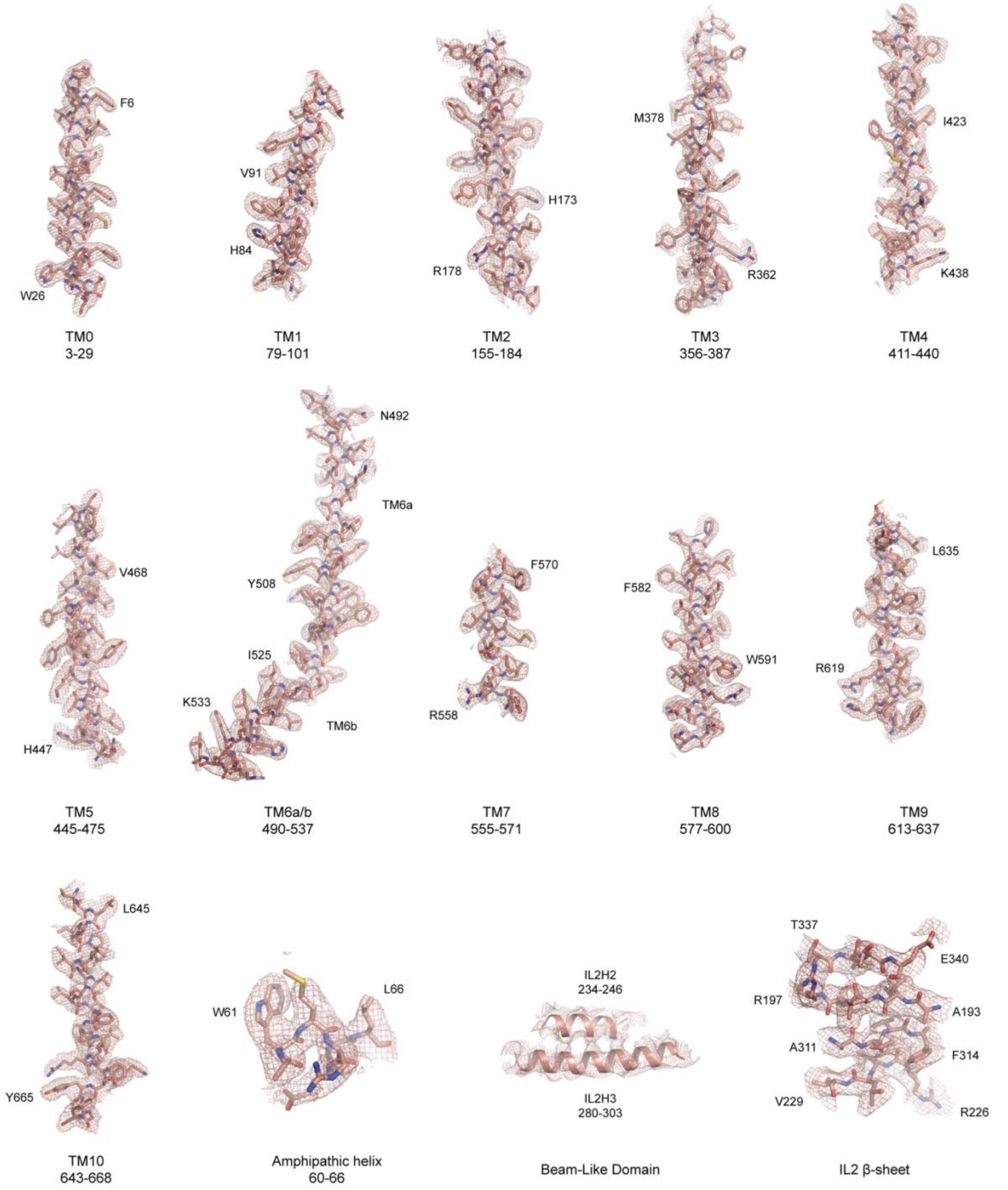


**Supplementary Figure 2: Fit of OSCA3.1 model to LocalDeblur map.**
Fit of OSCA3.1 model to selected regions of the LocalDeblur map. Map was contoured at a threshold of 2 (for the BLD) or 4 σ.

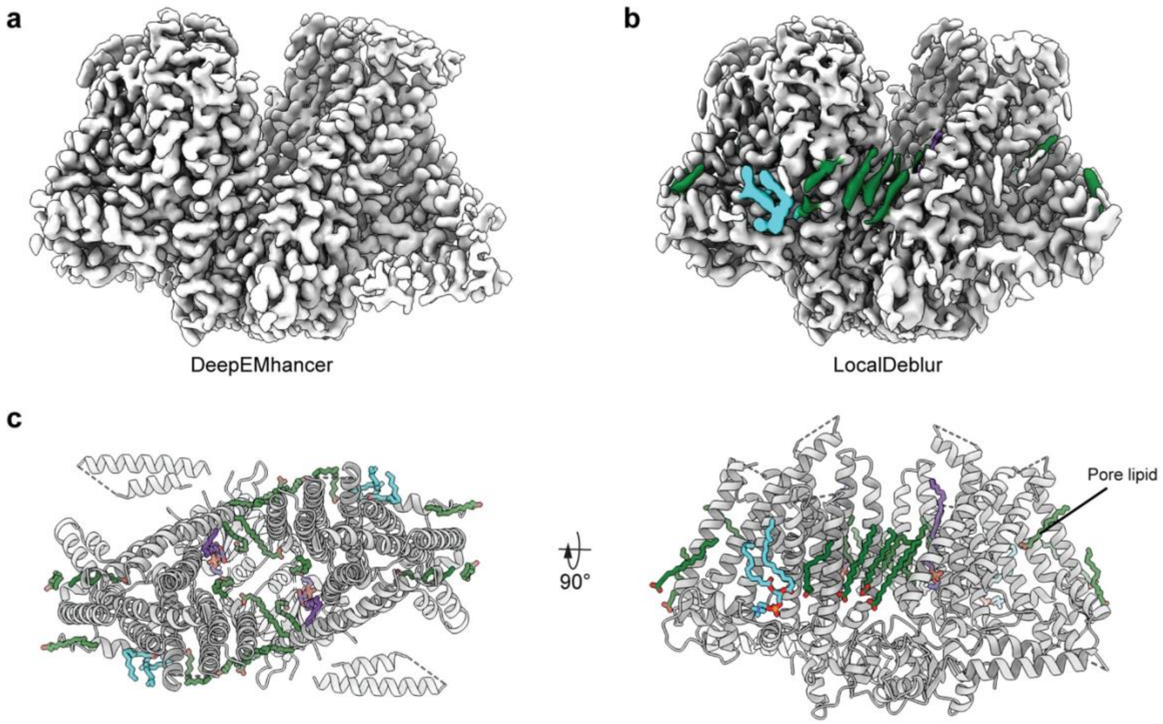


**Supplementary Figure 3: Comparison of DeepEMhancer and LocalDeblur postprocessed maps.**
**a,** DeepEMhancer map. **b,** LocalDeblur map with lipid-like densities colored. **c,** Top (left) and front (right) view of OSCA3.1 model with modelled lipids. Lipids were tentatively modelled as palmitic acid (green), phosphatidylcholine (cyan), and lyso-phosphatidylethanolamine (purple).

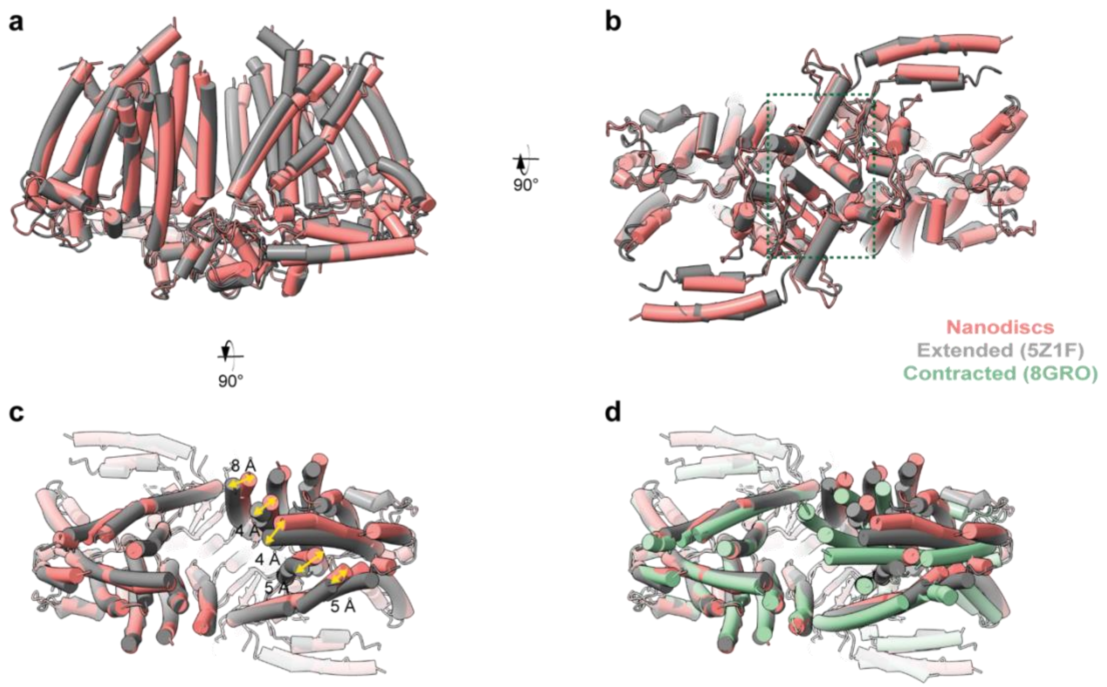


**Supplementary Figure 4: Comparison of OSCA3.1 in nanodiscs and detergent states.**

**a,** Front view of model superposition of OSCA3.1 in nanodiscs and extended state in digitonin (PDB: 5Z1F). Alignment done on left protomer. **b,** Intracellular view. Dimerization domain is enclosed in a green square. **c,** Extracellular view illustrating shifts of the nanodisc sample towards the outer edges, and increasing the distance between the two protomers in the inter-subunit cleft. **d,** Extracellular view of the OSCA3.1 in nanodiscs, extended state in digitonin, and contracted state in LMNG (PDB: 8GRO).

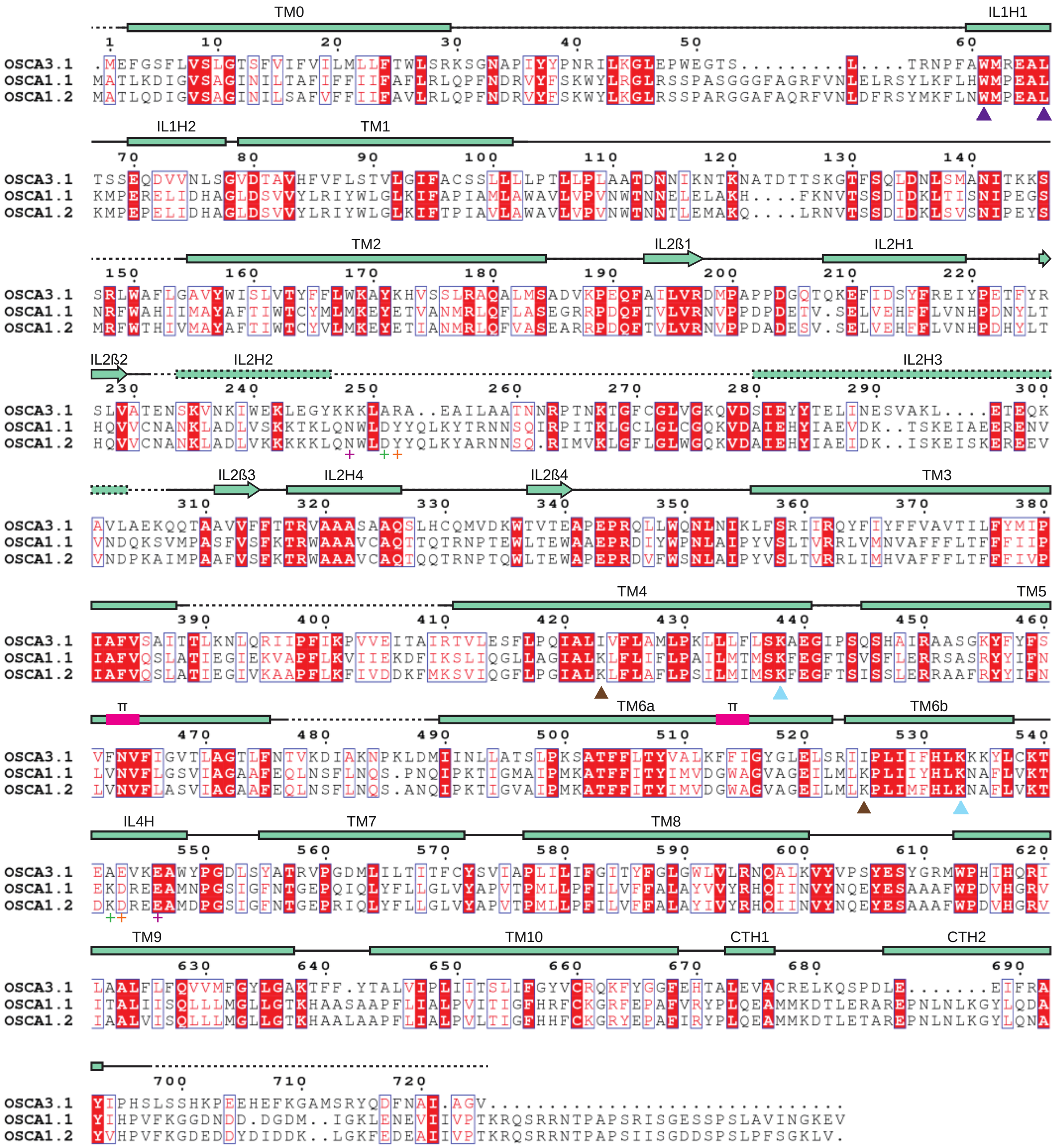

**Supplementary Figure 5: Protein sequence alignment of OSCA3.1.**
Amino acid sequence alignment of *Arabidopsis thaliana* OSCA3.1, OSCA1.1 and OSCA1.2. Secondary structure of OSCA3.1 represented on top, where rectangles represent α helices and arrows β strands. Dashed lines correspond to regions not modelled. π-helical turns labeled in magenta. IL2H2 and IL2H3 were modelled as poly-Ala and the registry was assigned tentatively. Purple arrowheads point to residues in the amphipathic helix that were mutated in OSCA1.2 for electrophysiology experiments. Predicted lipid-interacting residues denoted by light blue (conserved between OSCA1.2 and OSCA3.1) or brown (not conserved) arrowheads. Evolutionarily coupled residues

marked by plus sign (+), with each pair having the same color. TM: Transmembrane; IL: Intracellular Loop; CT: C-terminal; H: Helix; ß: ß-strand.

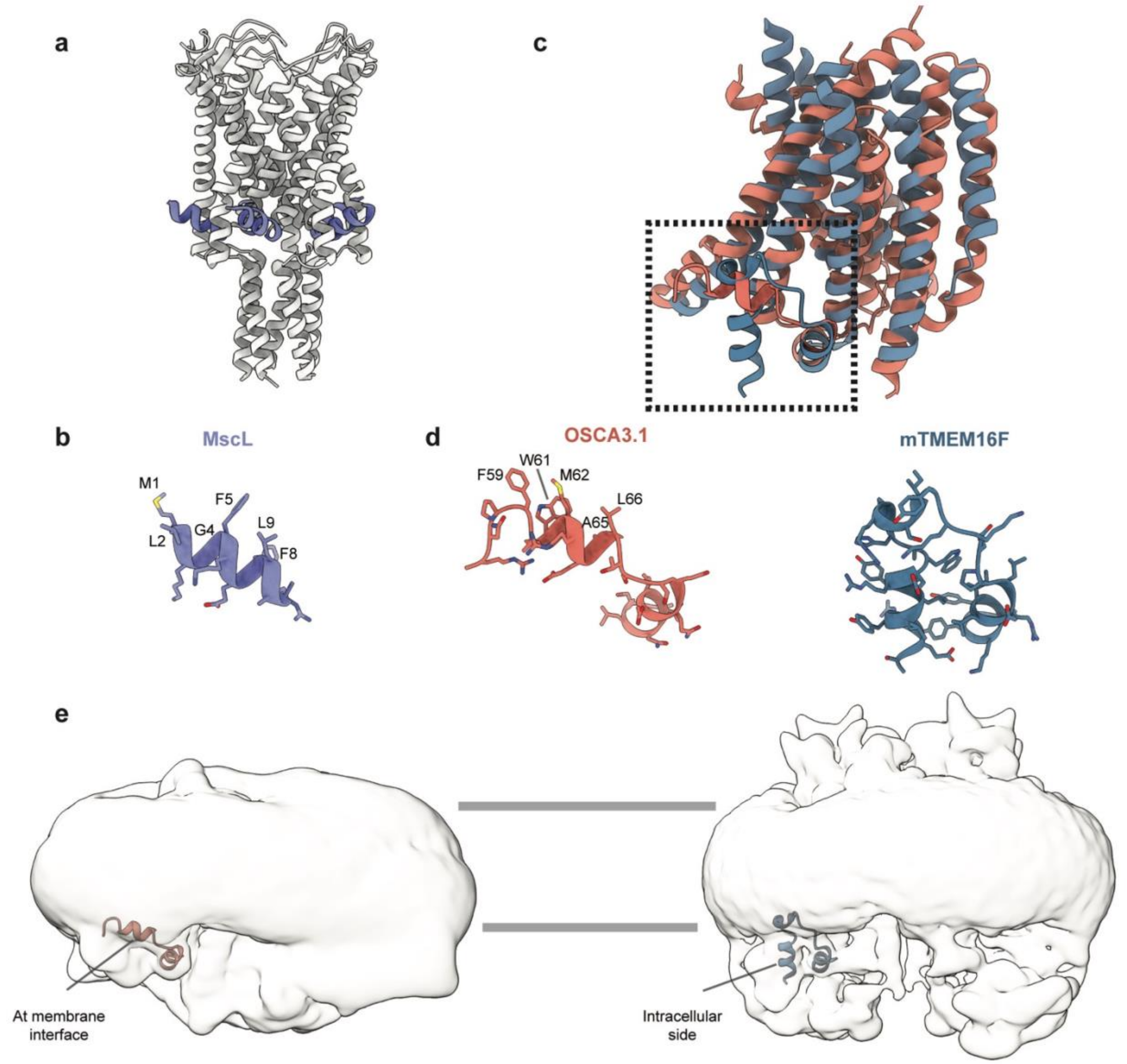


**Supplementary Figure 6: Comparison of the amphipathic helix of OSCA3.1 with corresponding helices in TMEM16 structures.**

**a,** Structure of MscL (PDB:2OAR). The amphipathic helix is highlighted in purple. **b,** Amphipathic helix of MscL with lipid-facing residues labeled. **c,** Superposition of transmembrane region of one protomer of OSCA3.1 (TM0 was excluded for simplicity) and homologous mouse (m) TMEM16F (PDB:6QP6). **d,** Inset from **c**: zoom-in view of amphipathic helix of OSCA3.1 (left) and homologous intracellular helix of mTMEM16F (right). Lipid-facing residues of OSCA3.1 are labeled. **c,** EM density of unsharpened density of OSCA3.1 (left) and mTMEM16F (right, EMDB: 4611) showing the position of respective helices shown in **c** and **d**. A gaussian-filter of 2 σ was applied to the densities. Horizontal gray lines denote the approximate position of the membrane bilayer.

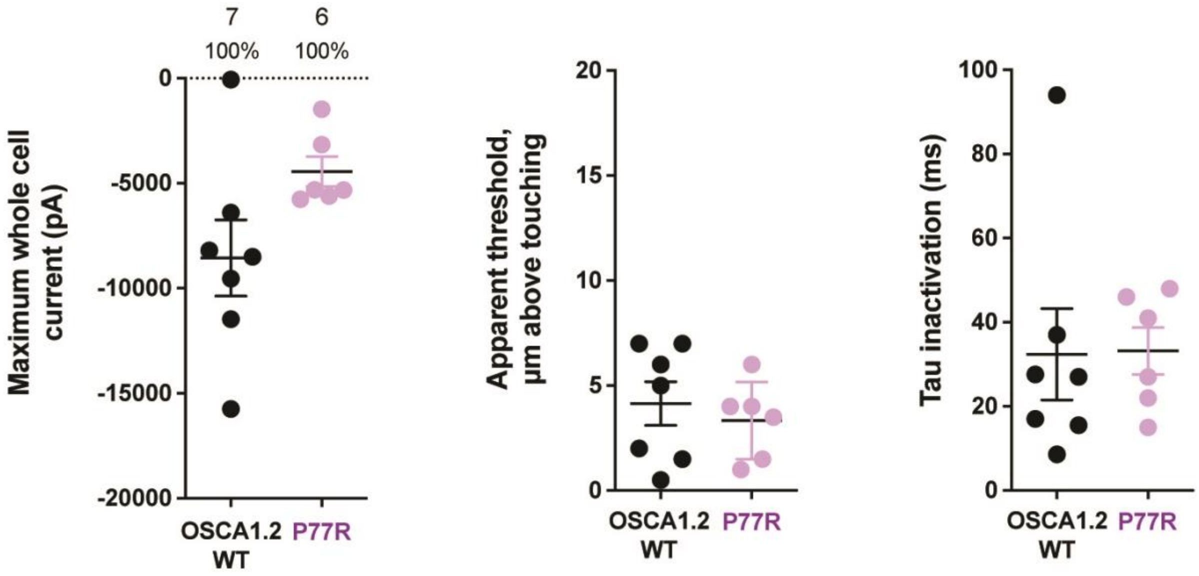


**Supplementary Figure 7: Poke-induced response of OSCA1.2$_{P77R}$.**
Introduction of arginine from OSCA3.1 into the OSCA1.2 background did not alter the poke-induced response. Left panel: Maximum poke-induced whole-cell currents observed for cells exposed to displacements up to 7.2 ± 1.4 µm (N=7; mean ± S.E.M.) and 6.5 ± 0.6 µm (N=6) above touching for WT and OSCA1.2$_{P77R}$, respectively. Center panel: the apparent threshold in µm above touching for the cells in this cohort. Right panel: Time constant of inactivation of currents elicited in the cells in this cohort.

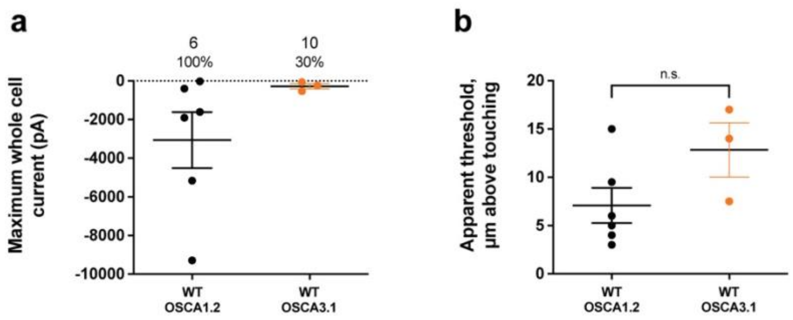


**Supplementary Figure 8: OSCA3.1 may be poke-sensitive at high thresholds.** Poke-induced whole-cell current responses are observed in HEK-P1KO cells expressing OSCA3.1. **a,** Maximum poke-induced currents observed in whole-cell mode for cells exposed to displacements up to 10.4 ± 1.2 µm (N=6; mean ± S.E.M.) and 13.8 ± 0.9 µm (N=3) above touching for $OSCA1.2_{WT}$ and $OSCA3.1_{WT}$, respectively. Although 6 of 6 OSCA1.2 expressing cells responded to poking, only 3 of 10 showed a response before the disruption of the recording. **b,** The apparent threshold in µm above touching the cell for this cohort.

**Supplementary Table 1. Data collection, processing, model refinement and validation.**

| | OSCA3.1 |
|---|---|
| **Data collection and processing** | |
| Magnification | 29000 |
| Voltage (kV) | 300 |
| Electron exposure (e–/Å$^2$) | 50 |
| Defocus range (μm) | -0.4 to -1.5 |
| Pixel size (Å) | 1.03 |
| Initial particle images (no.) | 1,913,316 |
| Symmetry imposed | C2 |
| Final particle images (no.) | 197,944 |
| Map resolution (Å) | 2.6 |
| FSC threshold | 0.143 |
| Map sharpening *B* factor (Å$^2$) | -64 |
| **Model** | |
| Composition | |
| Peptide chains | 2 |
| Protein residues | 1148 |
| Ligands | 24 |
| R.m.s. deviations | |
| Bond lengths (Å) | 0.023 |
| Bond angles (°) | 1.677 |
| Validation | |
| MolProbity score | 0.74 |
| Clashscore | 0.74 |
| EMRinger score | 4.12 |
| Poor rotamers (%) | 0.00 |
| Ramachandran plot | |
| Favored (%) | 98.93 |
| Allowed (%) | 1.07 |
| Disallowed (%) | 0.00 |
| **Deposition ID** | |
| EMDB | 41911 |
| PDB | 8U53 |
